# Complex-based Ligand-Binding Proteins Redesign by Equivariant Diffusion-based Generative Models

**DOI:** 10.1101/2024.04.17.589997

**Authors:** Viet Thanh Duy Nguyen, Nhan D. Nguyen, Truong Son Hy

**Author notes:** These authors contributed equally to this work (Viet Thanh Duy Nguyen, Nhan D. Nguyen). Correspondence to Truong Son Hy.

## Abstract

Proteins, serving as the fundamental architects of biological processes, interact with ligands to perform a myriad of functions essential for life. Designing functional ligand-binding proteins is pivotal for advancing drug development and enhancing therapeutic efficacy. In this study, we introduce ProteinReDiff, an efficient computational framework targeting the redesign of ligand-binding proteins. Using equivariant diffusion-based generative models, ProteinReDiff enables the creation of high-affinity ligand-binding proteins without the need for detailed structural information, leveraging instead the potential of initial protein sequences and ligand SMILES strings. Our evaluations across sequence diversity, structural preservation, and ligand binding affinity underscore ProteinReDiff’s potential to advance computational drug discovery and protein engineering. Our source code is publicly available at https://github.com/HySonLab/Protein_Redesign.

## Introduction

Proteins, often referred to as the molecular architects of life, play a critical role in virtually all biological processes. A significant portion of these functions involves interactions between proteins and ligands, underpinning the complex network of cellular activities. These interactions are not only pivotal for basic physiological processes, such as signal transduction and enzymatic catalysis, but also have broad implications in the development of therapeutic agents, diagnostic tools, and various biotechnological applications ^1–3^. Despite the paramount importance of protein-ligand interactions, the majority of existing studies have primarily focused on protein-centric designs to optimize specific protein properties, such as stability, expression levels, and specificity^4–8^. This prevalent approach, despite leading to numerous advancements, does not fully exploit the synergistic potential of optimizing both proteins and ligands for redesigning ligand-binding proteins. By embracing an integrated design approach, it becomes feasible to refine control over binding affinity and specificity, leading to applications such as tailored therapeutics with reduced side effects, highly sensitive diagnostic tools, efficient biocatalysis, targeted drug delivery systems, and sustainable bioremediation solutions^9–11^, thus illustrating the transformative impact of redesigning ligand-binding proteins across various fields.

Traditional methods for designing ligand-binding proteins have relied heavily on experimental techniques, characterized by systematic but often inefficient trial-and-error processes^12–14^. These methods, while foundational, are time-consuming, resource-intensive, and sometimes fall short in precision and efficiency. The emergence of computational design has marked a transformative shift, offering new pathways to accelerate the design process and gain deeper insights into the molecular basis of protein-ligand interactions. However, even with the advancements in computational approaches, significant challenges remain. Many existing models demand extensive structural information, such as protein crystal structures and specific binding pocket data, limiting their applicability, especially in urgent scenarios like the emergence of novel diseases ^15–17^. For instance, during the outbreak of a new disease like COVID-19, the spike proteins of the virus may not have well-characterized binding sites, delaying the development of effective drugs^18,19^. Furthermore, the complexity of binding mechanisms, including allosteric effects and cryptic pockets, adds another layer of difficulty^20,21^. Specifically, many proteins do not exhibit clear binding pockets until ligands are in close vicinity, necessitating extensive simulations to reveal potential binding interfaces^21,22^. While molecular dynamics simulations offer detailed atomistic insights into binding mechanisms, they often prove inadequate for designing high-throughput sequences due to high computational cost^9,23^. This complexity underscores the need for a drug design methodology that is agnostic to predefined binding pockets.

Our study addresses those identified challenges by introducing ProteinReDiff, a computational framework developed to enhance the process of redesigning ligand-binding proteins. Originating from the foundational concepts of the Equivariant Diffusion-Based Generative Model for Protein-Ligand Complexes (DPL)^24^, ProteinReDiff incorporates key improvements inspired by the representation learning modules from the AlphaFold2 (AF2) architecture ^25^. Specifically, we integrate the Outer Product Update (adapted from outer product mean of AF2), Single Representation Attention (adapted from MSA row attention module), and Triangle Multiplicative Update modules into our Residual Feature Update procedure. These modules collectively enhance the framework’s ability to capture intricate protein-ligand interactions, improve the fidelity of binding affinity predictions, and enable more precise redesigns of ligand-binding proteins.

The framework integrates the generation of diverse protein sequences with blind docking capabilities. Starting with a selected protein-ligand pair, our approach stochastically masks amino acids and equivariantly denoises the diffusion model to capture the joint distribution of ligand and protein complex conformations. Another key feature of our method is blind docking, which predicts how the redesigned protein interacts with its ligand without the need for predefined binding site information, while relying solely on initial protein sequences and ligand SMILES strings^26^. This streamlined approach significantly reduces reliance on detailed structural data, thus expanding the scope for sequence-based exploration of proteinligand interactions.

In summary, the contributions of our paper are outlined as follows:

- We introduce ProteinReDiff, an efficient computational framework for ligand-binding protein redesign, rooted in equivariant diffusion-based generative models. Our innovation lies in integrating AF2’s representational learning modules to enhance the framework’s ability to capture intricate protein-ligand interactions.
- Our framework enables the design of high-affinity ligand-binding proteins without reliance on detailed structural information, relying solely on initial protein sequences and ligand SMILES strings.
- We comprehensively evaluate our model’s outcomes across multiple design aspects, including sequence diversity, structure preservation, and ligand binding affinity, ensuring a holistic assessment of its effectiveness and applicability in various contexts.

## Related Work

### Traditional Approaches in Protein Design

Protein design has historically hinged on computational and experimental strategies that paved the way for modern advancements in the field. These foundational methodologies emphasized the intricate balance between understanding protein structure and engineering novel functionalities, albeit with inherent limitations in scalability and precision. Key traditional approaches include:

- **Rational Design**^27–29^ focused on introducing specific mutations into proteins based on known structural and functional insights. This method required an in-depth understanding of the target protein structures and how changes might impact its function.
- **Directed Evolution** ^30–33^ mimicked natural selection in the laboratory, evolving proteins towards desired traits through iterative rounds of mutation and selection. Despite its effectiveness in discovering functional proteins, the process was often labor-intensive and time-consuming.

These traditional methods have been instrumental in advancing our understanding and capability in protein design. However, their limitations in terms of efficiency, specificity, and the broad applicability of findings highlighted the need for more versatile and scalable approaches. As the field progressed, the integration of computational power and biological understanding opened new avenues for innovation in protein design, leading to the exploration and adoption of more advanced methodologies.

### Deep Generative Models in Protein Design

Since their inception, deep generative models have significantly advanced fields like computer vision (CV)^34^ and natural language processing (NLP)^35^, sparking interest in their application to protein design. This enthusiasm has led to numerous studies that harness these models for innovating within the protein design area. Among these, certain types of deep generative models have distinguished themselves through their effectiveness and the promising results they have achieved, including:

- **Variational Autoencoders (VAEs)** are harnessed for their ability to learn rich representations of protein sequences, enabling the generation of novel sequences through manipulation in the latent space^36–38^.
- **Autoregressive models** predict the probability of each amino acid in a sequential manner, facilitating the generation of coherent and functionally plausible protein sequences^39,40^.
- **Generative Adversarial Networks (GANs)** employ two networks that work in tandem to produce protein sequences indistinguishable from real ones, enhancing the realism and diversity of generated designs^41,42^.
- **Diffusion models** represent a step forward by gradually transforming noise into structured data, simulating the complex process of folding sequences into functional proteins^43–46^.

However, the majority of these studies have focused on protein-centric designs, with a noticeable gap in research that integrates both proteins and ligands for the purpose of redesigning ligand-binding proteins. Such integration is crucial for a holistic understanding of the intricate dynamics between protein structures and their ligands, a domain that remains underexplored.

### Current Approaches in Ligand-Binding Protein Redesign

#### Heavy Reliance on Detailed Structural Information

Contemporary computational methodologies for designing proteins that target specific surfaces predominantly rely on structural insights from native complexes, underscoring the critical role of fine-tuning side-chain interactions and optimizing backbone configurations for optimal binding affinity ^15–17,44,47,48^. These strategies often initiate with the generation of protein backbones, employing inverse folding techniques to identify sequences capable of folding into these pre-designed structures^6,7,48,49^. This approach signifies a paradigm shift by prioritizing structural prediction ahead of sequence identification, aiming to produce proteins that not only fit the desired conformations for potential ligand interactions but also navigate around the challenge of undefined binding sites. Despite the advantages, including the potential of computational docking to create binders via manipulation of antibody scaffolds and varied loop geometries ^36,50,51^, a notable challenge persists in validating these binding modes with high-resolution structural evidence. Additionally, the traditional focus on a limited array of hotspot residues for guiding protein scaffold placement often restricts the exploration of possible interaction modes, particularly in cases where target proteins lack clear pockets or clefts for ligand accommodation^22,52^.

#### Limited Training Data and Lack of Diversity

Existing approaches often rely on a limited set of training data, which can restrict the diversity and generalizability of the resulting models. For instance, datasets like PDBBind provide detailed ligand information, but their scope is limited^53^. This limitation is further compounded when protein datasets lack corresponding ligand data, reducing the effectiveness of the training process. Traditional methodologies also tend to focus on a narrow range of protein-ligand interactions, potentially overlooking the broader spectrum of possible interactions.

#### Single-Domain Denoising Focus

Previous methodologies typically concentrate on denoising either in sequence space or structural space, but not both. Approaches like ProteinMPNN^6^, LigandMPNN^17^, and MIF^48^ primarily operate in sequence space, while others like DPL function in structural space^24^. This single-domain focus can limit the ability to capture the full complexity of protein-ligand interactions, which inherently involve both sequence and structural dimensions. Consequently, these methodologies may fall short of accurately predicting the functional capabilities of redesigned proteins.

#### Challenges in Generating Diverse Sequences with Structural Integrity

While some approaches prioritize sequence similarity to generate functional proteins, they often do so at the expense of structural integrity. For example, ProteinMPNN and CARP focus heavily on sequence similarity, which can result in a lack of diversity and flexibility in the generated sequences^6,7^. This limitation can hinder the ability to explore a wider range of functional conformations, reducing the effectiveness of the protein design process.

#### Distinct Improvements of Our Approach

We address the weaknesses of available methodologies by integrating diverse datasets, employing a dual-domain denoising strategy, and ensuring the generation of diverse sequences while maintaining structural integrity. Our approach utilizes only protein sequences and ligand SMILES strings, eliminating the need for detailed structural information. By combining PDBBind^53^ and CATH^54^ datasets, we effectively double our training data, enhancing protein representations. Our equivariant and KL-divergence loss functions enable denoising across both sequence and structural dimensions, capturing the full complexity of protein-ligand interactions. This approach maintains structural fidelity and promotes sequence diversity, overcoming the limitations of methodologies prioritizing sequence similarity at the expense of diversity.

## Background

### Protein Language Models (PLMs)

Protein Language Models (PLMs) harness the power of natural language processing (NLP) to unravel the intricate latency embedded within protein sequences. By analogizing amino acid sequences to human language sentences, PLMs unlock profound insights into protein functions, interactions, and evolutionary trajectories^55^. These models leverage advanced text processing techniques to predict structural, functional, and interactional properties of proteins based solely on their amino acid sequences^56–59^. Their adoption in protein design has catalyzed significant progress, with studies leveraging PLMs to translate protein sequence data^47,60–62^ into actionable insights, thus guiding the precise engineering of proteins with targeted functional attributes.

Mathematically, a PLM can be represented as a function *F* that maps a sequence of amino acids *S* = [*s*_1_, *s*_2_, …, *s*_*n*_], where *s*_*i*_ denotes the *i*-th amino acid in the sequence, to a high-dimensional feature space that encapsulates the protein’s structural and functional properties:

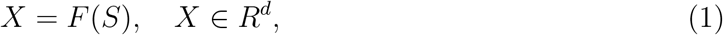

where *X* represents the continuous representation or embedding derived from the sequence *S* and *d* represents the dimensionality of the embedding space, determined by the PLM’s architecture. This embedding captures the complex dependencies and patterns underlying the protein’s structural information and biological functionality. Through training on known sequences and structures, PLMs discern the “grammar” governing protein folding and function, facilitating accurate predictions.

We employ the ESM-2 model^59^, a state-of-the-art protein language model with 650 million parameters, pre-trained on nearly 65 million unique protein sequences from the UniRef^63^ database, to feature initial masked protein sequences. ESM-2 enriches the latent representation of protein sequences, bypassing the need for conventional multiple sequence alignment (MSA) methods. By incorporating structural and evolutionary information from input sequences, ESM-2 enables us to unravel interaction patterns across protein families for effective ligand targeting. This understanding is crucial for designing and optimizing ligand-binding proteins.

### Equivariant Diffusion-based Generative Models

We utilize a generative model driven by equivariant diffusion principles, drawing from the foundations laid by Variational Diffusion Models^64^ and E(3) Equivariant Diffusion Models^65^.

#### The Diffusion Procedure

First, we employ a diffusion procedure that is equivariant with respect to the coordinates of atoms *x*, alongside a series of progressively more perturbed versions of *x*, known as latent variables *z*_*t*_, with *t* varying from 0 to 1. To maintain translational invariance within the distributions, we opt for distributions on a linear subspace that anchors the centroid of the molecular structure at the origin, and designate *N*_*x*_ as a Gaussian distribution within this specific subspace. The conditional distribution of the latent variable *z*_*t*_ given *x*, for any given *t* in the interval [0, 1], is defined as

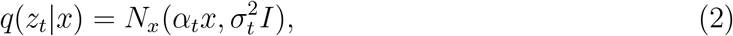

where *α*_*t*_ and 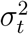 represent strictly positive scalar functions of *t*, dictating the extent of signal preservation versus noise introduction, respectively. We implement a variance-conserving mechanism where 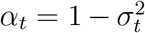 and posit that *α*_*t*_ smoothly and monotonically decreases with *t*, ensuring *α*_0_ ≈ 1 and *α*_1_ ≈ 0. Given the Markov property of this diffusion process, it can be described via transition distributions as

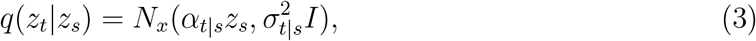

for any *t* > *s*, where *α*_*t*|*s*_ = *α*_*t*_/*α*_*s*_ and 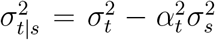. The Gaussian posterior of these transitions, conditional on *x*, can be derived using Bayes’ theorem:

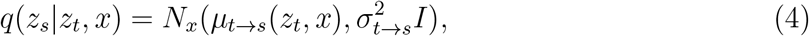

With

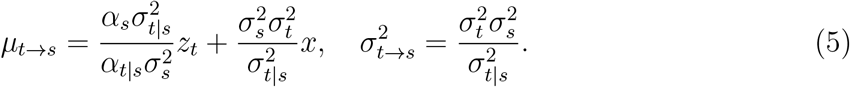

#### The Generative Denoising Process

The construction of the generative model inversely mirrors the diffusion process, generating a reverse temporal sequence of latent variables *z*_*t*_ from *t* = 1 back to *t* = 0. By dividing time into *T* equal intervals, the generative framework can be described as:

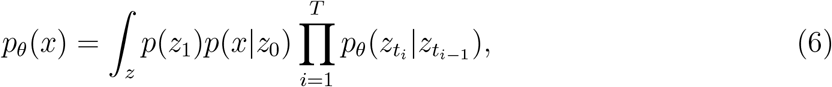

with *s*(*i*) = (*i* − 1)/*T* and *t*(*i*) = *i*/*T*. Leveraging the variance-conserving nature and the premise that *α*_1_ ≈ 0, we posit *q*(*z*_1_) = *N*_*x*_(0, *I*), hence treating the initial distribution of *z*_1_ as a standard Gaussian:

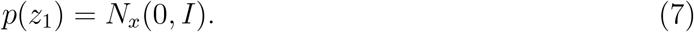

Furthermore, under the variance-preserving framework and assuming *α*_0_ ≈ 1, the distribution *q*(*z*_0_ | *x*) is modeled as highly peaked^64,66^. This allows us to approximate *p*_data_(*x*) as nearly constant within this narrow peak region. This yields:

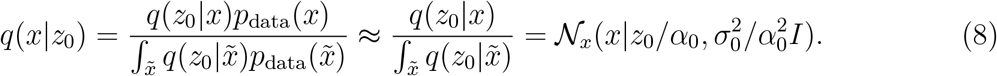

Accordingly, we approximate *q*(*x*|*z*_0_) through:

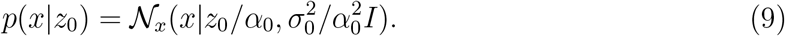

The generative model’s conditional distributions are then formulated as:

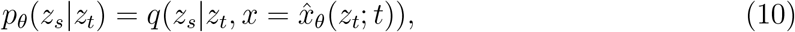

which mirrors *q*(*z*_*s*_|*Fsz*_*t*_, *x*) but substitutes the actual coordinates *x* with the estimates from a temporal denoising model 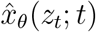, which employs a neural network parameterized by *θ* to predict *x* from its noisier version *z*_*t*_. This denoising model’s framework, predicated on noise prediction 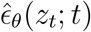, is articulated as:

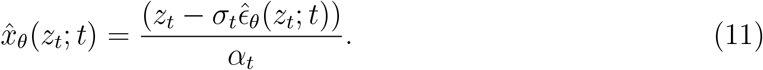

Consequently, the transition mean 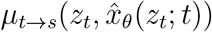 is determined by:

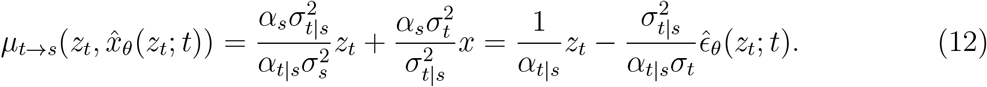

## Method

In this section, we detail the methodology employed in our noise prediction model, which is depicted in Figure 1 and consists of three main procedures: (1) input featurization, (2) residual feature update, and (3) equivariant denoising. Through these steps, we transform raw protein and ligand data into structured representations, iteratively refine their features, and leverage denoising techniques inherent in the diffusion model to improve sampling quality.

**Figure 1:**
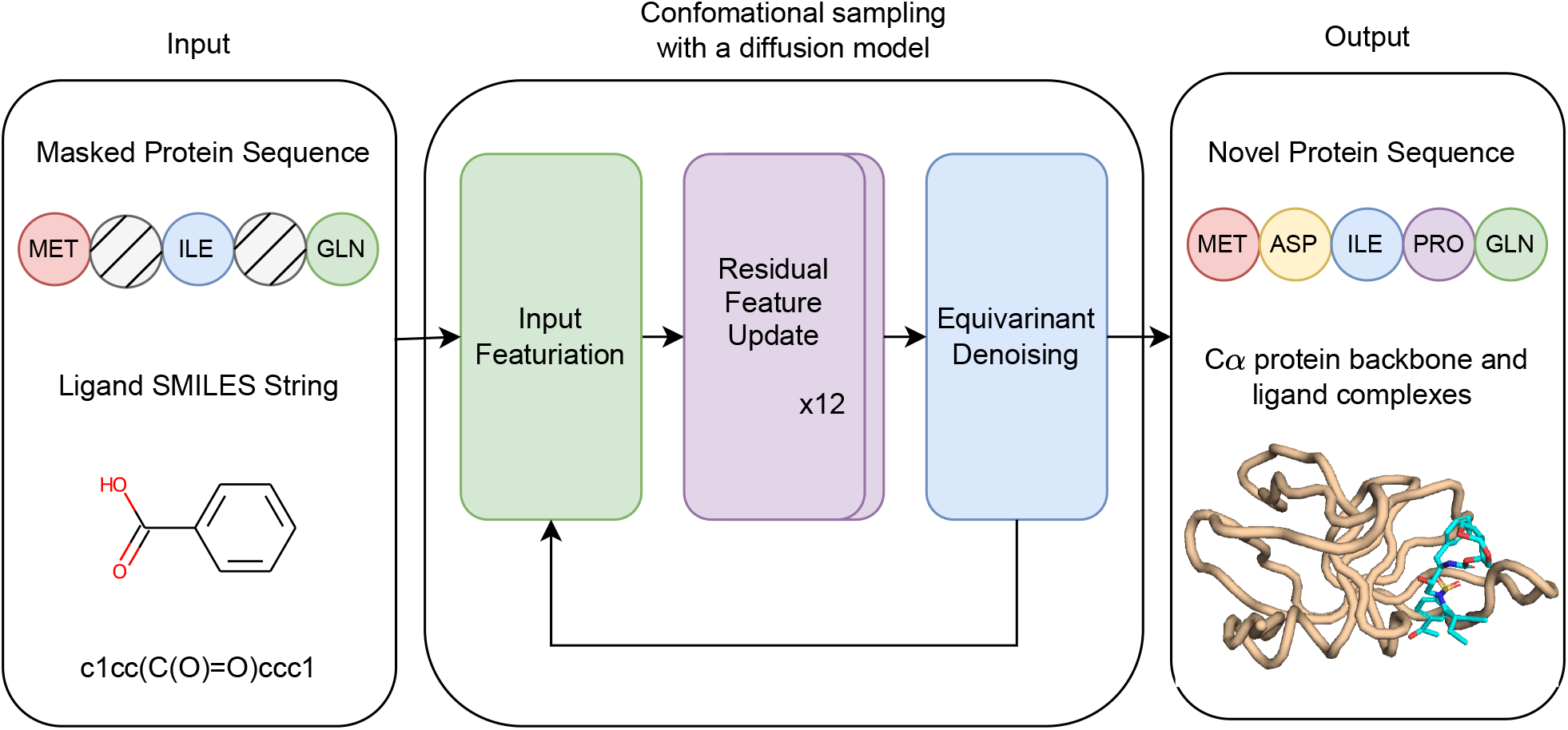
Overview of the proposed framework. The process begins with utilizing a protein amino acid sequence and a ligand SMILES string as inputs. The conformational sampling process includes iteratively applying input featurization, updating residual features, and denoising equivariantly, ultimately yielding novel protein sequences alongside their corresponding *Cα* protein backbone and ligand complexes.

### Input Featurization

We develop both single and pair representations from protein sequences and ligand SMILES string (Figure 2). For proteins, we initially applied stochastic masking to segments of the amino acid sequences. The protein representation is attained through the normalization and linear mapping of the output from the final layer of the ESM-2 model, which is subsequently combined with the amino acid and masked token embeddings. Additionally, for pair representations of proteins, we leveraged pairwise relative positional encoding techniques, drawing from established methodologies^25^. For ligand representations, we employed a comprehensive feature embedding approach, capturing atomic and bond properties such as atomic number, chirality, connectivity, formal charge, hydrogen attachment count, radical electron count, hybridization status, aromaticity, and ring presence for atoms; and bond type, stereochemistry, and conjugation status for bonds. These representations are subsequently merged, incorporating radial basis function embeddings of atomic distances and sinusoidal embeddings of diffusion times. Together, these steps culminate in the formation of preliminary complex representations, laying the foundation for our computational analyses.

**Figure 2:**
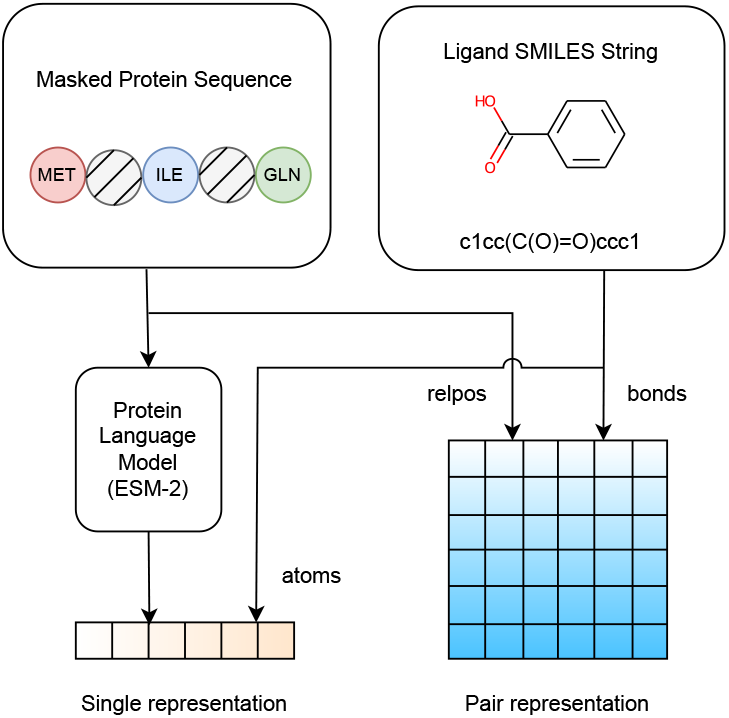
Overview of the input featurization procedure of the model.

**Figure 3:**
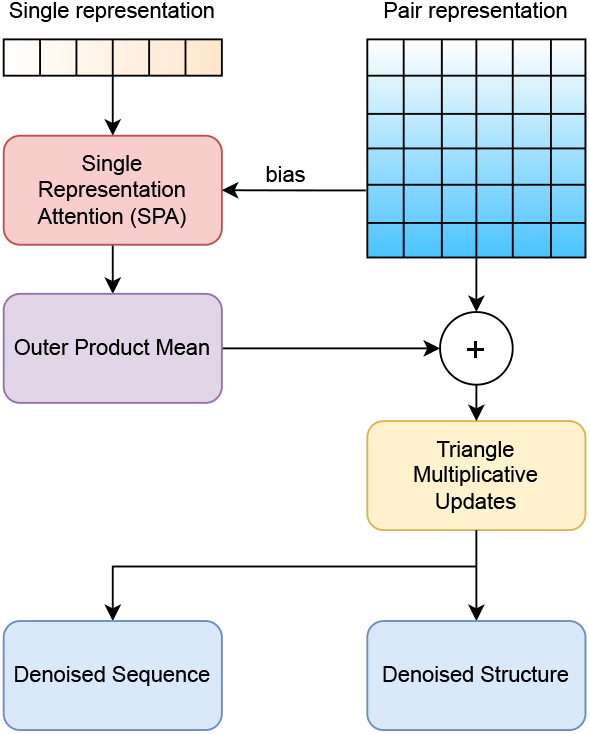
Overview of the residual feature update procedure of the model.

### Residual Feature Update Procedure

Our approach deviates significantly from the residual feature update procedure employed in the original DPL model^24^. While the DPL model relied on Alphafold2’s Triangular Multiplicative Update for updating single and pair representations, where these representations mutually influence each other, our objective is to optimize this procedure for greater efficiency. Specifically, we incorporate enhancements such as the Outer Product Update and Single Representation Attention to formulate sequence representational hypotheses of pro-tein structures and to model suitable motifs for binding target ligands specifically. These modules, integral to Evoformer, the sequence-based module of AF2, play a crucial role in extracting essential connections among internal motifs that serve structural functions (i.e., ligand binding) when structural information is not explicitly provided during training. Importantly, we adapt and tailor these modules to fit within our model architecture, ensuring their effectiveness in capturing the intricate interplay between proteins and ligands.

### Single Representation Attention Module

The Single Representation Attention (SRA) module, derived from the Alphafold2 model’s MSA row attention with pair bias, accounts for long-range interactions among residues and ligand atoms within a single protein-ligand embedding vector. In essence, the attention mechanism assigns importance to those involved in complex-based folding to denoise the equivariant loss (Section) in a self-supervised manner. While the original Alphafold2 MSA row attention mechanism processes input for a single sequence, the SRA module is designed to incorporate representations from multiple protein-ligand complexes concurrently. Specifically, the pair bias component of the SRA attention module captures dependencies between proteins and ligands, which was shown to fit the attention score better than the regular selfattention model without bias terms^67^. By considering both the single representation vector (which encodes the protein/ligand sequential representation) and the pairwise representation vector (which encodes protein-protein and protein-ligand interactions), this cross-attention mechanism exchanges information between pairwise and single representation to effectively preserves internal motifs, as evidenced by contact overlap metrics^55,68^. As transformer architecture is widely used for predicting protein functions^69^, we observed similar efficacy to our binding affinity prediction in section Results and Appendix **??**-**??**. For a detailed description of the computational steps implemented in this module, refer to Algorithm 1.

#### Algorithm 1

Single Representation Attention pseudocode

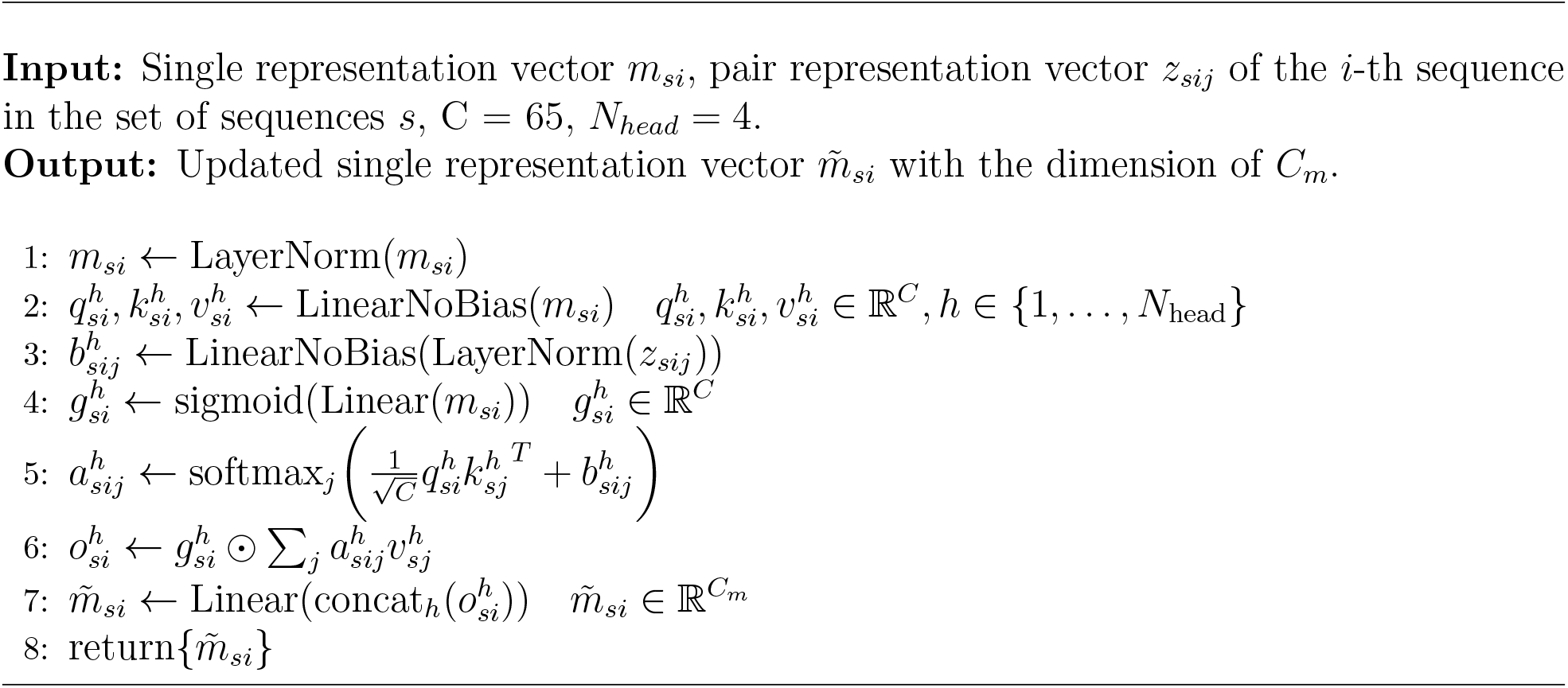

### Outer Product Update

Since the SRA encodings have a shape (*s, r, c*_*m*_) and the pair representation has a shape (*s, r, r, c*_*z*_), the outer product (OPU) layer merges insights by reshaping SRA encodings into pair representations. This module leverages evolutionary cues from ESM to generate plausible structural hypotheses for pair representations^70^. It first calculates the outer product of the SRA embeddings of protein-ligand pairs, then aggregates the outer products to yield a measure of co-evolution between every residue pair^55^. Analogous to Tensor Product Representations (TPR) in NLP, the outer product is akin to the filler-and-role binding relationship, where each entity (i.e. amino acid residue) on a sequence is attached to a rich functional embedding based on its relationship to one another^71–73^.

This process integrates correlated information of residues *i* and *j* of a sequence *s*, resulting in the intermediate Kronecker product tensors (.i.e. role embeddings in NLP) ^67,74,75^. Subsequently, an affine transformation projects those representations to hypotheses concerning the relative positions of residues *i* and *j* under biophysical constraints. Our implementation adapts the outer product without computing the mean to maintain the pair representations of multiple protein-ligand complexes. For a detailed description of the computational steps implemented in this module, refer to Algorithm 2.

#### Algorithm 2

Outer product update pseudocode

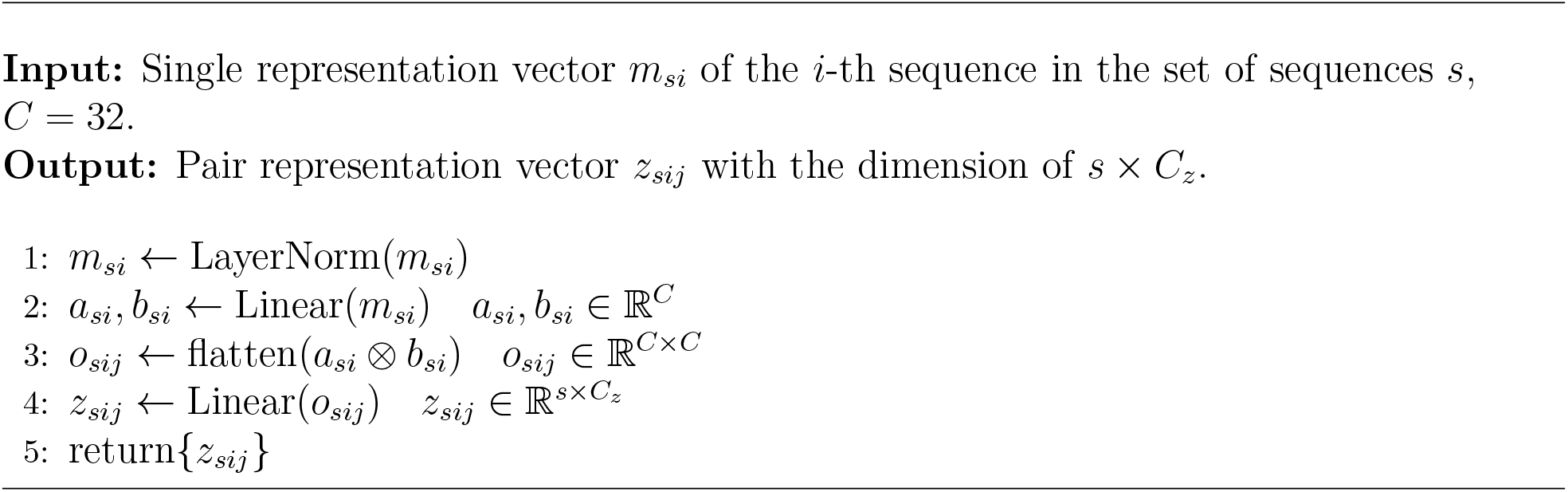

### Triangle Multiplicative Updates

After refining the pair representation, our model interprets the primary protein-ligand structure using principles from graph theory, treating each residue as a distinct entity interconnected through the pairwise matrix. These connections are then refined through triangular multiplicative updates to account for physical and geometric constraints, such as triangular inequality. While the SRA weights the importance of residues, the triangular multiplicative update acts as another stack of transformer-based layers where any two edges affect the third one to enforce triangle equivariance ^55,76^. The starting and ending nodes propagate information in and out of neighbors in similar fashion as the message-passing framework^67^. These mechanisms enable the model to generate more accurate representations of protein-ligand complexes, leading to improved predictive performance in predicting binding affinities and structural characteristics.

### Equivariant Denoising

During the equivariant denoising process, the final pair representation undergoes symmetrization and is then transformed using a multi-layer perceptron (MLP) into a weight matrix *W*. This matrix is utilized to compute the weighted sum of all relative differences in 3D space for each atom, as shown in the equation^24^:

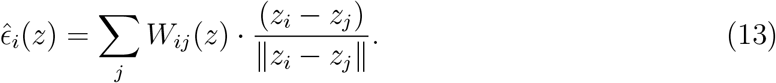

Afterward, the centroid is subtracted from this computation, resulting in the output of our noise prediction model 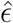. Additionally, it’s important to note that the described model maintains SE(3)-equivariance, meaning that:

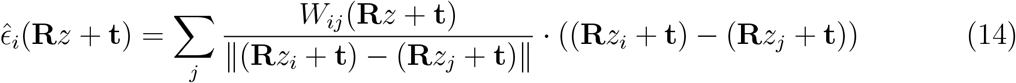

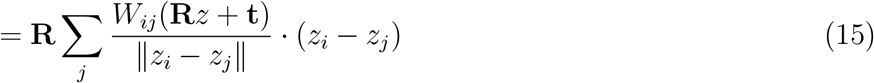

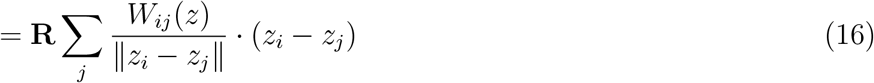

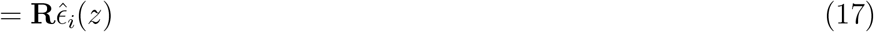

for any rotation **R** and translation **t**. This property is derived from the fact that the final representation, and hence the weight matrix *W*, depends solely on atom distances that are invariant to rotation and translation.

## Experiments

### Training Process

#### Materials

Our training strategy leverages a meticulously curated dataset encompassing a broad range of protein structures, including both ligand-bound (holo) and ligand-free (apo) forms, sourced from two key repositories: PDBBind v2020^53^ and CATH 4.2^54^. PDBBind v2020 offers a diverse collection of protein-ligand complexes, while CATH 4.2 provides a substantial repository of protein structures. Each dataset was selected for its unique contributions to our understanding of protein-ligand interactions and structural diversity. This strategic selection of datasets ensures our model is exposed to a wide and varied spectrum of protein-ligand interactions and structural configurations, enabling comprehensive evaluation against diverse inverse folding benchmarks. By training on both holo and apo structures, our approach imbues the model with a robust understanding of protein-ligand dynamics, equipping it to navigate the complexities of unseen protein-ligand interaction scenarios effectively.

To ensure robust model training and evaluation, we employ careful data partitioning techniques. Using MMseqs2^77^, we clustered and partitioned the protein sets for training, validation, and testing, maintaining sequence similarities between 40% and 50% to ensure unbiased training and predictions, following protocols from other protein models^25,48^. For ligands, we cluster based on the Tanimoto similarity of Morgan fingerprints^78^ on ligand structures. Incorporating CATH 4.2 data into PDBBind not only preserves the objectivity of the train/test/validation partitions but also substantially decreases the similarities within ligand sets, as shown in Table 1.

**Table 1:** Similarity between Train/Validation/Test Sets of Proteins and Ligands. The values represent similarity percentages for the original PDBBind dataset versus combined PDBBind with CATH datasets in parentheses.

| **Protein** | Validation | Test | **Ligand** | Validation | Test |
| --- | --- | --- | --- | --- | --- |
| Train | 36.0% (36.2%) | 38.0% (42.2%) | Train | 72.2% (36.1%) | 9.41% (3.11%) |
| Validation | - | 39.08% (43.5%) | Validation | - | 9.37% (3.17%) |

Table 2 provides an overview of the partitioning details, facilitating a clear understanding of the distribution of samples across different subsets of the dataset.

**Table 2:** Data Partitioning Overview (Unit: number of samples)

| **Dataset** | **Train** | **Validation** | **Test** |
| --- | --- | --- | --- |
| PDBBind v2020 | 9430 | 552 | 207 |
| CATH 4.2 | 15261 | 939 | - |

- **PDBBind v2020**: For consistency and comparability with previous studies, we first adhered to the test/training/validation split settings outlined in established literature^79^, specifically following the configurations defined in the respective sources for the PDBBind v2020 datasets^80^. Then, we filtered out those highly similar sequences (above 95%) to keep the average similarities between 40%-50%.
- **CATH 4.2**: In our approach, we deliberately focused on proteins with fewer than 400 amino acids and less similar (below 90%) sequences from the CATH 4.2 database. This selective criterion was chosen to prioritize smaller proteins, which often represent more druggable targets of interest in drug discovery and development endeavors. During both the training and validation phases, SMILES strings of CATH 4.2 proteins were represented as asterisks (masked tokens) to denote unspecified ligands. Notably, CATH 4.2 was excluded from the test set due to the absence of corresponding ligands required for evaluating protein-ligand interactions.

### Loss Functions

Previous models typically denoise in only one domain, such as ProteinMPNN ^6^, LigandMPNN^17^, and MIF^48^ in sequence space, and DPL^24^ in structural space. This limitation restricts their ability to fully capture the intricate interactions between proteins and ligands. To address this, we have introduced significant modifications to the loss function to better suit the task of ligand-binding protein redesign. By tailoring the loss function to integrate both sequence and structural spaces, our approach effectively addresses the unique challenges of proteinligand interactions. Specifically, the optimization of our model for ligand-binding protein redesign is governed by a composite loss function *L*, formulated as follows:

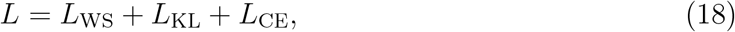

#### Weighted Sum of Relative Differences (*L*_WS_)

This component ensures the model’s sensitivity to the directional influence between atoms, supporting the accurate prediction of the denoised structure while maintaining physical symmetries. It is crucial for the equivariant denoising step, enabling accurate noise prediction for atoms in the protein-ligand complex. The loss is defined as:

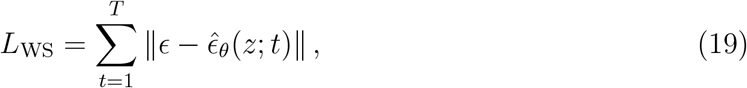

where *T* is the total number of time steps in the diffusion process, *ϵ* is the Gaussian noise vector 𝒩(**0, I**), and 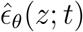 is the loss prediction at time step *t* parameterized by a weight MLP in Section.

#### Kullback-Leibler Divergence (*L*_KL_) ^81^

This component quantifies the divergence between the model’s predictions and actual sequence data at timestep *t* − 1, playing a pivotal role in the denoising process. Defined as *KL*(*x*_pred t-1_, *seq*_t-1_), it contrasts the predicted distribution, *x*_pred t-1_, against the true sequence distribution, *seq*_t-1_, leveraging the diffusion process’s *γ* parameter for temporal adjustment. This loss is also applied in the Protein Generator^5^ model to ensure the model’s predictions progressively align with actual data distributions, enhancing the accuracy of sequence and structure generation by minimizing the expected divergence.

#### Cross-entropy Loss (*L*_CE_)

This loss function is crucial for the accurate prediction of protein sequences, aligning them with the ground truth through effective classification. It denoises each amino acid from masked latent embedding to a specific class, leveraging categorical cross-entropy to rigorously penalize discrepancies between the model’s predicted probability distributions and the actual distributions for each amino acid type.

### Training Performance

Throughout the training phase, we meticulously observed the model’s performance, paying close attention to the dynamics between training and validation losses, as demonstrated in Figure 4. While the training loss consistently diminished, indicating effective learning, the validation loss exhibited more variability. Despite these fluctuations, the validation loss showed an overall downward trend, suggesting that the model is improving its generalization capabilities over time. The general alignment between the downward trends of training and validation losses indicates that the model is learning effectively without significant overfitting.

**Figure 4:**
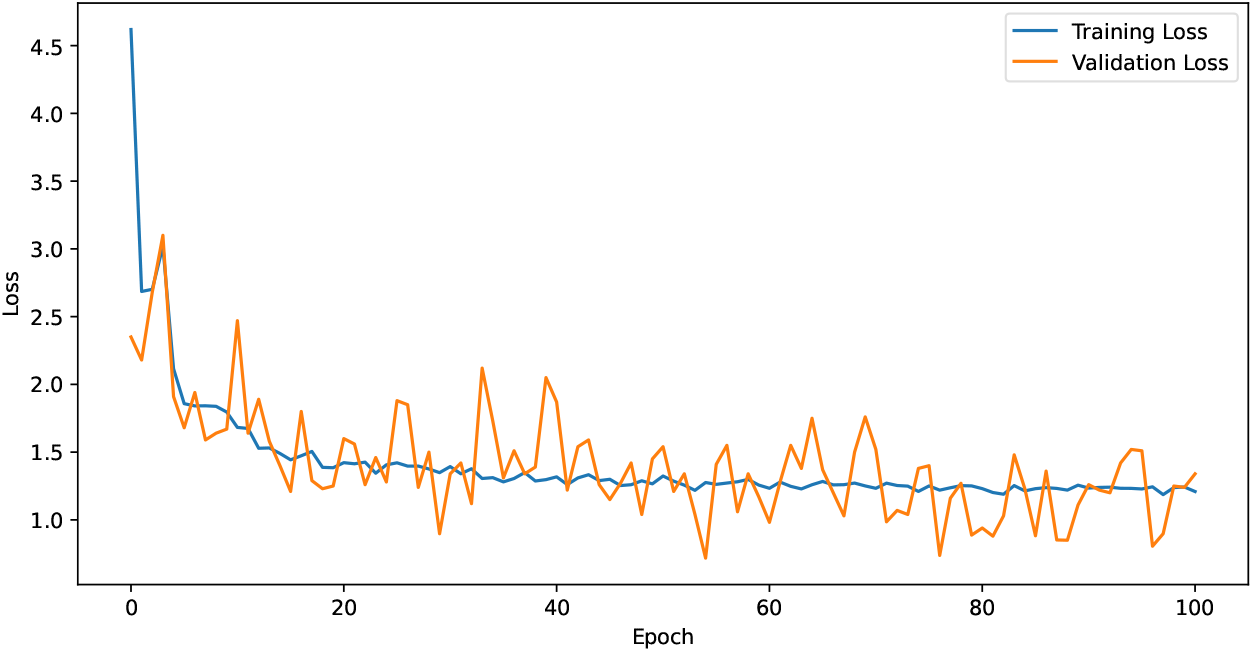
Training history chart of ProteinReDiff, showcasing the evolution of training and validation losses over epochs.

### Evaluation Process

#### Ligand Binding Affinity (LBA)

Ligand binding affinity is a fundamental measure that quantifies the strength of the interaction between a protein and a ligand. This metric is crucial as it directly influences the effectiveness and specificity of potential therapeutic agents; higher affinity often translates to increased drug efficacy and lower chances of side effects^82^. Within this context, ProteinReDiff is evaluated on its ability to generate protein sequences for significantly improved binding affinity with specific ligands. We utilize a docking score-based approach for this assessment, where the docking score serves as a quantitative indicator of affinity. Expressed in kcal/mol, these scores inversely relate to binding strength — lower scores denote stronger, more desirable binding interactions.

#### Sequence Diversity

Sequence diversity is crucial for exploring protein’s functional space^83^. It reflects the capacity of our model, ProteinReDiff, to traverse the vast landscape of protein sequences and generate a wide array of variations. To quantitatively assess this diversity, we utilize the average edit distance (Levenshtein distance)^84^ between all pairs of sequences generated by the model. This metric offers a nuanced measure of variability, surpassing traditional metrics that may overlook subtle yet significant differences. The diversity score is calculated using the formula:

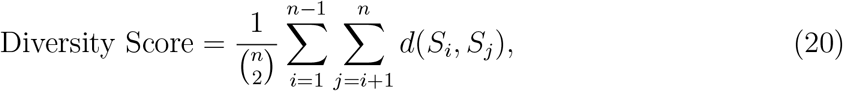

where *d*(*S*_*i*_, *S*_*j*_) represents the edit distance between any two sequences *S*_*i*_ and *S*_*j*_. This calculation provides an empirical gauge of ProteinReDiff’s ability to enrich the protein sequence space with novel and diverse sequences, underlining the practical variance introduced by our model.

#### Structure Preservation

Structural preservation is paramount in the redesign of proteins, ensuring that essential functional and structural characteristics are maintained post-modification. To effectively measure structural preservation between the original and redesigned proteins, three key metrics: the Template Modeling Score (TM Score)^85^, the Root Mean Square Deviation (RMSD)^86^, and the Contact Overlap (CO)^87^. These two metrics collectively provide a comprehensive assessment of structural integrity and similarity, essential for evaluating the success of our protein redesign efforts.

**The Root Mean Square Deviation (RMSD)** is a measure used to quantify the distance between two sets of points. In the context of protein structures, these points are the positions of the atoms in the protein. The RMSD is given by the formula:

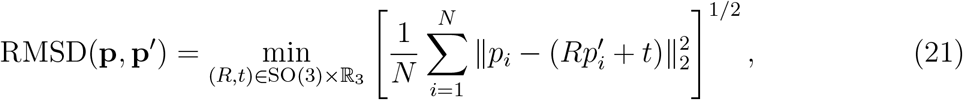

Where 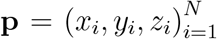 and 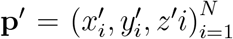 denote two sequences of *N* 3D coordinates representing the atomic positions in the original and redesigned proteins, respectively. This formula calculates the minimum root mean square of distances between corresponding atoms, after optimal superposition, which involves finding the best-fit rotation *R* and translation *t* that aligns the two sets of points. A lower RMSD value indicates a higher degree of structural similarity, making it a direct measure of the extent to which structural deviation has been minimized. Achieving a low RMSD is desirable, as it signifies that the redesign process has successfully preserved the core structural configuration of the original protein.

**TM Score** provides a normalized measure of structural similarity between protein configurations, which is less sensitive to local variations and more reflective of the overall topology. The TM Score is defined as follows:

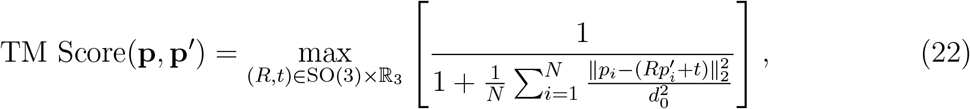

where *d*_0_ is a scale parameter typically chosen based on the size of the proteins. The closer the TM Score is to 1, the more similar the structures are, indicating global structural alignment.

**Contact Overlap (CO)** provides a complementary perspective to RMSD and TM Score by focusing on the preservation of local structural motifs rather than overall geometric similarity. Several studies show that having high CO indicates protein’s residue pairs having co-evolutionary signals^87,88^ and performing related functions^89^. CO quantitatively measures the conservation of inter-atomic contacts between the original and redesigned protein structures, which are crucial for the protein’s structural integrity and functional capabilities. The metric is defined as:

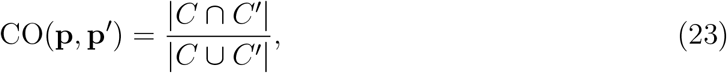

where *C* = *{*(*i, j*) : ∥*p*_*i*_ − *p*_*j*_∥ < *r*_*c*_, *i*≠ *j}* and 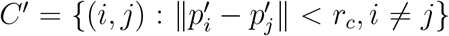 represent the sets of contacts in the original and redesigned proteins, respectively. Here, *p*_*i*_ and 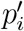 are the positions of atoms in the original and redesigned proteins, and *r*_*c*_ is a predefined cutoff distance that determines when two atoms are considered to be in contact. A high CO score indicates that many of the original contacts are preserved in the redesigned structure, suggesting that the redesign maintains much of the original protein’s structural network, which is crucial for its stability and function.

#### Experimental Setup

To evaluate ProteinReDiff, we employed Omegafold^90^ to predict the three-dimensional structures of all designed protein sequences. The choice of Omegafold over AF2 was favorable because Omegafold can more accurately fold proteins with low similarity to existing proteomes, making it suitable for proteins lacking available ligand-binding conformations. Next, we utilized AutoDock Vina^91^ to conduct docking simulations and evaluate the binding affinity between the redesigned proteins and their respective ligands based on the predicted 3D structures. To ensure fair comparisons and mitigate potential biases introduced by predocked structures, we aligned our redesigned protein structures with reference structures before docking. This approach is crucial, particularly because the use of pre-docked structures may favor certain conformations, leading to inaccurate evaluations. Additionally, to provide context for our results, we compared the binding scores of our redesigned proteins not only with those of the original proteins but also with proteins generated by other protein design models. Although these models may exhibit different sequence characteristics compared to those explicitly designed for ligand binding affinity, comparing their scores offers valuable insights. Such comparisons help elucidate the interplay between protein sequence and structure in determining ligand interactions, enriching the interpretation of our findings and advancing our understanding of protein-ligand interactions.

#### Benchmark Model Selection

In selecting benchmark models for performance comparison, we focused on state-of-the-art approaches, particularly those relevant to protein design tasks. Traditionally, protein design has been primarily based on inverse folding, utilizing protein structure information. Our choices encompass a range of methodologies:

- MIF^48^, MIF-ST^48^, and ProteinMPNN^6^ are notable for generating sequences with high identity and experimental significance, utilizing protein structure information.
- The Protein Generator^5^, a representative of RosettaFold models^44^, employs diffusion-based methods, making it an intriguing comparative candidate. The model also shares a similar loss function, *L*_*KL*_, in sequence space with our model but diverges in modules and training procedures (i.e., stochastic masking).
- ESMIF^49^, belonging to the ESM model family^92^, stands as another competitive benchmark, emphasizing the generation of high-quality sequences.
- CARP, while lacking ligand information, shares similar protein input and output characteristics with our models, warranting inclusion for comparison.
- DPL^24^, originally geared towards protein-ligand complex generation, was adapted for our purposes by modifying loss functions and incorporating a sequence prediction module, given its alignment with our model architecture.
- LigandMPNN^17^, resembling the most to our task in designing ligand-binding proteins, necessitates binding pocket information, unlike our model, which emphasizes a simplified yet effective approach for ligand-binding protein tasks.

Our model’s design prioritizes simplicity in input while achieving effectiveness in output for ligand-binding protein tasks. For a comprehensive comparison of input-output dynamics across each model, please consult Table 3.

**Table 3:** Comparison of protein design models based on input and output characteristics.

| Model | Input |  |  |  | Output |  |  |
| --- | --- | --- | --- | --- | --- | --- | --- |
|  | Protein Sequence | Protein Structure | Ligand SMILES | Binding Pocket | Protein Sequence | Protein Structure | Ligand Structure |
| CARP<sup>7</sup> | ✓ | × | × | × | ✓ | × | × |
| ESMIF<sup>49</sup> | × | ✓ | × | × | ✓ | × | × |
| MIF<sup>48</sup> | ✓ | ✓ | × | × | ✓ | × | × |
| MIF-ST<sup>48</sup> | ✓ | ✓ | × | × | ✓ | × | × |
| ProteinMPNN<sup>6</sup> | × | ✓ | × | × | ✓ | × | × |
| LigandMPNN<sup>17</sup> | × | ✓ | ✓ | ✓ | ✓ | × | × |
| Protein Generator<sup>5</sup> | × | ✓ | × | × | ✓ | × | × |
| DPL<sup>24</sup> | ✓ | × | ✓ | × | × | ✓ | ✓ |
| ProteinReDiff (Ours) | ✓ | × | ✓ | × | ✓ | ✓ | ✓ |

### Results and Discussion

We conducted comprehensive evaluation of ProteinReDiff, as detailed in Table 4 and visually represented in Figure 6, across the metrics of ligand binding affinity, sequence diversity, and structure preservation. These evaluations provide a clear depiction of the model’s performance relative to established baselines and within its variations.

**Table 4:** Comparison of method performance across multiple metrics: Ligand binding affinity (LBA), sequence diversity, and structure preservation. Ligand binding affinity (LBA), TM Score, and RMSD are reported as mean values with their respective margins of error.

| Category | Method | LBA<br>(kcal/mol) ↓ | Sequence<br>diversity ↑ | Structure preservation |  |  |
| --- | --- | --- | --- | --- | --- | --- |
|  |  |  |  | TM Score ↑ | RMSD (Å) ↓ | CO ↑ |
| Baseline | CARP | -5.658 ± 0.301 | 185.532 | 0.850 ± 0.023 | 3.768 ± 0.553 | 0.922 ± 0.003 |
|  | MIF | -5.518 ± 0.381 | 185.600 | **0.877** ± 0.020 | **2.986** ± 0.468 | 0.938 ± 0.002 |
|  | MIF-ST | -5.596 ± 0.330 | 185.584 | 0.872 ± 0.021 | 3.026 ± 0.451 | 0.937 ± 0.003 |
|  | ESMIF | -5.555 ± 0.326 | 187.512 | 0.837 ± 0.021 | 4.000 ± 0.501 | 0.915 ± 0.003 |
|  | ProteinMPNN | -5.423 ± 0.225 | 188.792 | 0.714 ± 0.026 | 6.806 ± 0.616 | 0.859 ± 0.004 |
|  | LigandMPNN | -5.717 ± 0.287 | **191.384** | 0.782 ± 0.024 | 4.512 ± 0.668 | 0.915 ± 0.008 |
|  | Protein Generator | -5.674 ± 0.266 | 186.962 | 0.806 ± 0.022 | 4.431 ± 0.523 | 0.899 ± 0.003 |
|  | DPL | -5.551 ± 0.459 | 188.139 | 0.788 ± 0.024 | 5.094 ± 0.537 | 0.896 ± 0.009 |
|  | Reference cases | -5.847 ± 0.263 | - | - | - | - |
| ProteinReDiff<br>(Ours) | 5% Masking | -5.805 ± 0.252 | 185.935 | 0.864 ± 0.022 | 3.197 ± 0.470 | **0.942** ± 0.007 |
|  | 15% Masking | **-6.803** ± 0.329 | 186.627 | 0.845 ± 0.023 | 3.690 ± 0.508 | 0.935 ± 0.007 |
|  | 30% Masking | -5.769 ± 0.244 | 187.877 | 0.803 ± 0.024 | 4.467 ± 0.544 | 0.916 ± 0.008 |
|  | 40% Masking | -5.617 ± 0.366 | 188.600 | 0.756 ± 0.026 | 5.639 ± 0.625 | 0.896 ± 0.008 |
|  | 60% Masking | -5.467 ± 0.318 | 190.425 | 0.305 ± 0.024 | 18.056 ± 0.773 | 0.735 ± 0.010 |
|  | 70% Masking | -5.470 ± 0.199 | 187.291 | 0.147 ± 0.004 | 23.197 ± 0.497 | 0.689 ± 0.007 |

For ProteinReDiff, we aimed to capture the diverse conformations of ligand-binding proteins, recognizing that they can adopt multiple structural states. To assess these conformations, we employed alignment metrics such as TM score, RMSD, and contact overlap (CO). In Figure 5, we presented several instances where the contact overlap appeared to be maintained, yet the RMSD is large and TM score is low. This discrepancy suggests that while global alignment metrics like TM score and RMSD may not adequately capture the domain shift within these complex ensembles, the preservation of local motifs, as indicated by contact overlap, remains crucial in our framework. This underscores the importance of capturing both global and local structural features for a comprehensive understanding of protein-ligand interactions.

**Figure 5:**
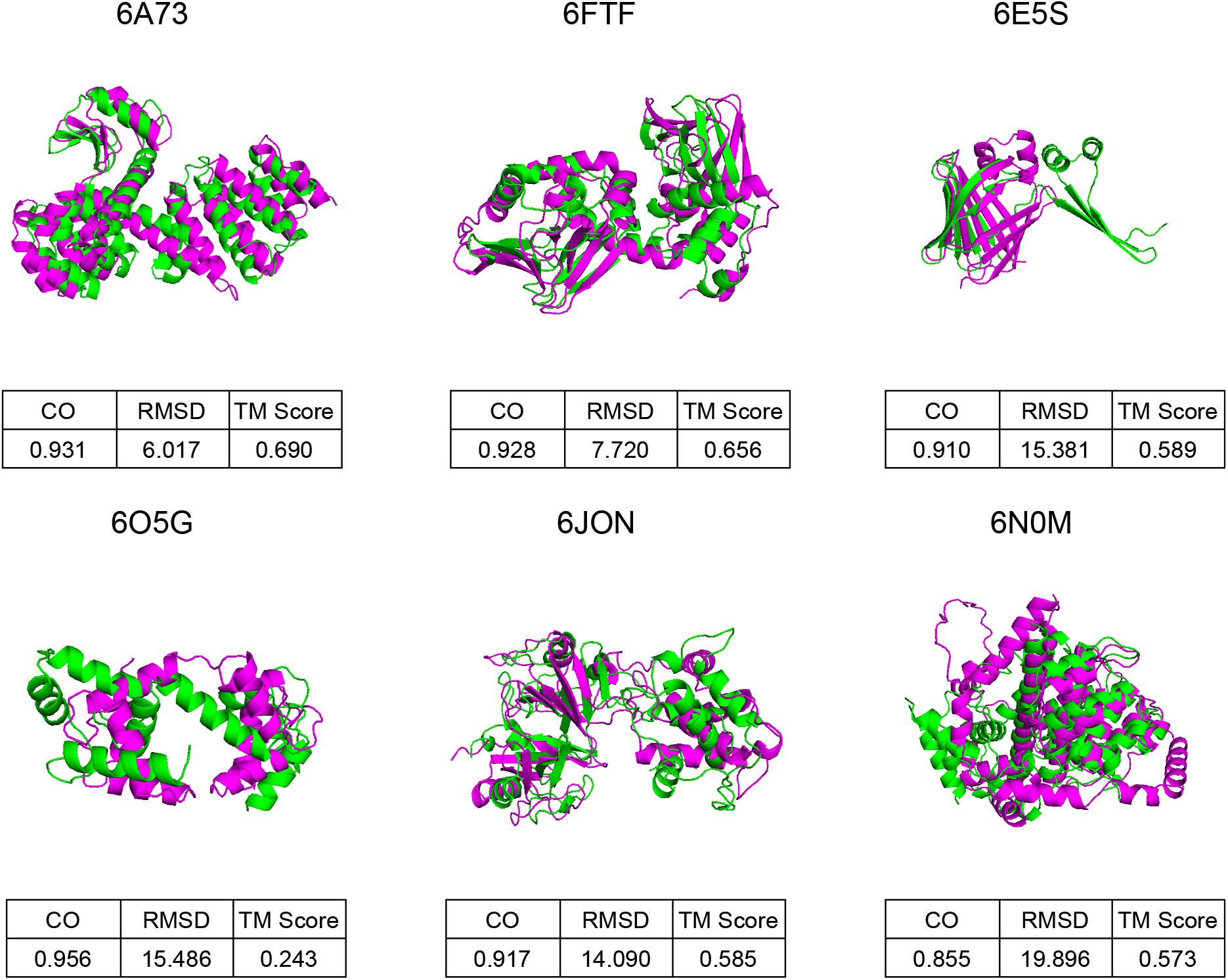
Comparative visualizations of protein structures, each annotated with its corresponding PDB ID. The figure includes a succinct table detailing Contact Overlap (CO) and Root Mean Square Deviation (RMSD) metrics. Original protein structures are highlighted in green, and the redesigned versions by ProteinReDiff are depicted in pink, illustrating the precise structural changes and enhancements achieved through the redesign.

**Figure 6:**
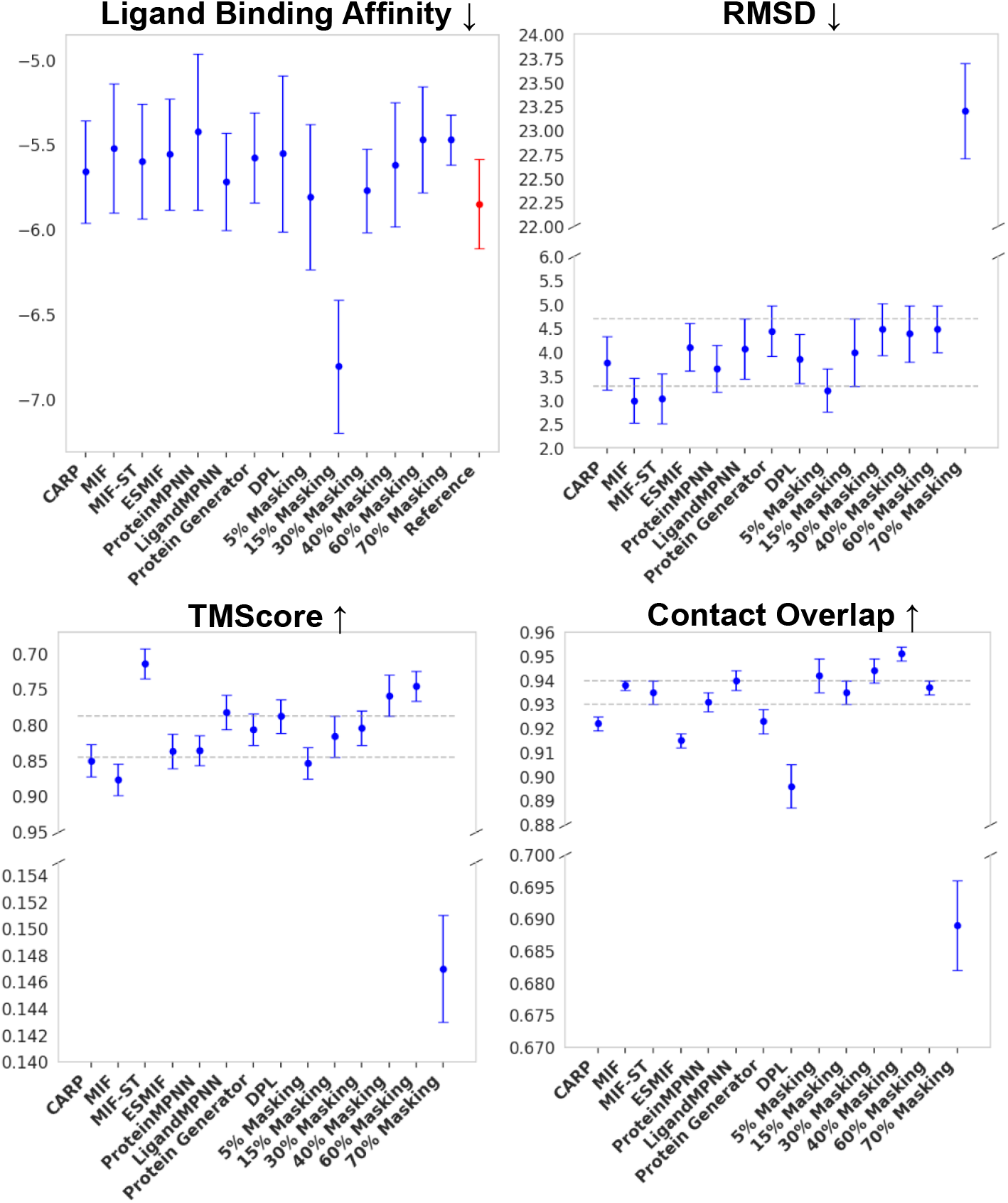
Visualization of method performance across metrics. The metrics are plotted with mean values and margins of error. For LBA, the red bar (top right) shows the docking score of reference complexes. The horizontal dash lines indicate the regions of 15% masking model which is our standard for comparison.

A pivotal observation from our study is ProteinReDiff’s unparalleled ability to enhance ligand binding affinity, particularly at a 15% masking ratio in Figure 6. This configuration not only surpasses the performance of Inverse Folding (IF) models and the original DPL framework but also exceeds the binding efficiencies of the original protein designs. By incorporating attention modules from AlphaFold2, ProteinReDiff effectively captures the complex interplay between proteins and ligands, demonstrating its superiority over the original DPL model. While other masking ratios within ProteinReDiff show varying degrees of effectiveness, lower ratios, though at the same par as reference, do not achieve the peak LBA performance observed at 15%. For instance, the 5% masked model emphasizes structural consistency with a high TM-Score and low RMSD, but does not exhibit the same level of binding capability as the 15% masking. These findings are also consistent with ablation studies shown in Appendix **??**. Conversely, higher masking ratios fail to strike the necessary balance between introducing beneficial modifications and maintaining functional precision, underscoring the importance of optimizing the masking ratio.

Our analysis of sequence diversity and structure preservation metrics reveals a delicate balance essential in protein redesign. The 15% masking ratio, identified as optimal for enhancing ligand binding affinity in our model, also aligns closely with benchmark methods in both sequence diversity and structure preservation. For instance, LigandMPNN excels in sequence diversity but faces challenges in obtaining binding pocket inputs for various design tasks, unlike our approach. Moreover, our models (at 30% and 40% maskings) significantly outperform others in contact overlap, crucial for diversifying structures while preserving functional motifs in protein redesign tasks. This equilibrium underscores ProteinReDiff’s ability to optimize ligand interactions without compromising the exploration of sequence diversity or the integrity of original protein structures.

In contrast, extreme values in either sequence diversity or structure preservation, which could be seen in other masking ratios, do not lead to optimal ligand binding affinities. This finding highlights an inverse relationship between pushing the limits of diversity and preservation and achieving the primary goal of binding enhancement. Thus, the 15% masking ratio not only stands out for its ability to significantly improve ligand binding affinity but also for maintaining a balanced approach, ensuring that enhancements in functionality do not detract from the protein’s structural and functional viability.

In Figure 7, we compare the ligand-binding affinity (LBA) of original and redesigned proteins by ProteinReDiff. The redesigned proteins maintain their original folds while significantly enhancing LBA. In ablation studies (Section), we can apply various masking strategies to adjust both sequence diversity and structural integrity. This approach has potential applications in different settings to control the affinity of ligand binders.

**Figure 7:**
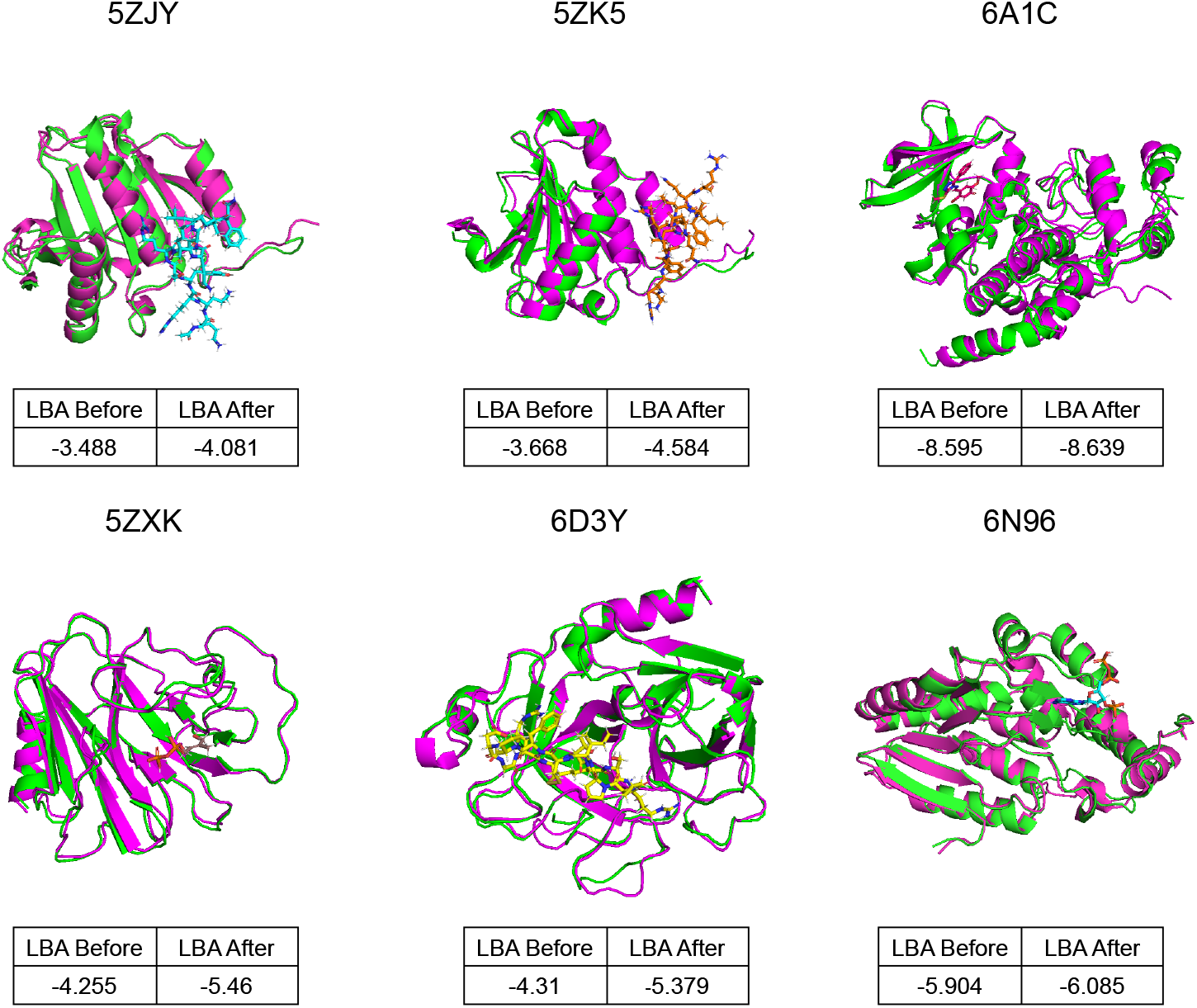
Comparative visualizations of protein-ligand complexes, each labeled with corresponding PDB IDs and accompanied by a small table showing Ligand Binding Affinity (LBA) before and after the redesign. Original structures are highlighted in green, while redesigned versions by ProteinReDiff appear in pink. Ligands are depicted in various colors to emphasize specific binding sites and molecular interaction enhancements post-redesign.

### Ablation Studies

Here we conducted thorough ablation studies on ProteinReDiff’s model architecture, featurization, and masking ratios. For complete ablation setup, please refer to Table **??** (Appendix **??**)

#### Interpreting Model Architecture

We trained ablated versions of ProteinReDiff without the SRA or OPU modules and compared them to the original DPL model. Initially designed for generating ensembles of complex structures, DPL was adapted for targeted protein redesign by adding sequence-based loss functions to generate new target sequences.

In Figure 8, we computed the performance score by averaging the sum of five evaluation metrics introduced in Sections,, and. Since the sequence diversity is not within the [0,1] range, we applied Min-Max normalization. For LBA and RMSD, we used inverse normalization to ensure that a score closer to 1.0 indicates better model performance. The average score is then compared with the baseline score of ProteinReDiff which was trained without any ablations.

**Figure 8:**
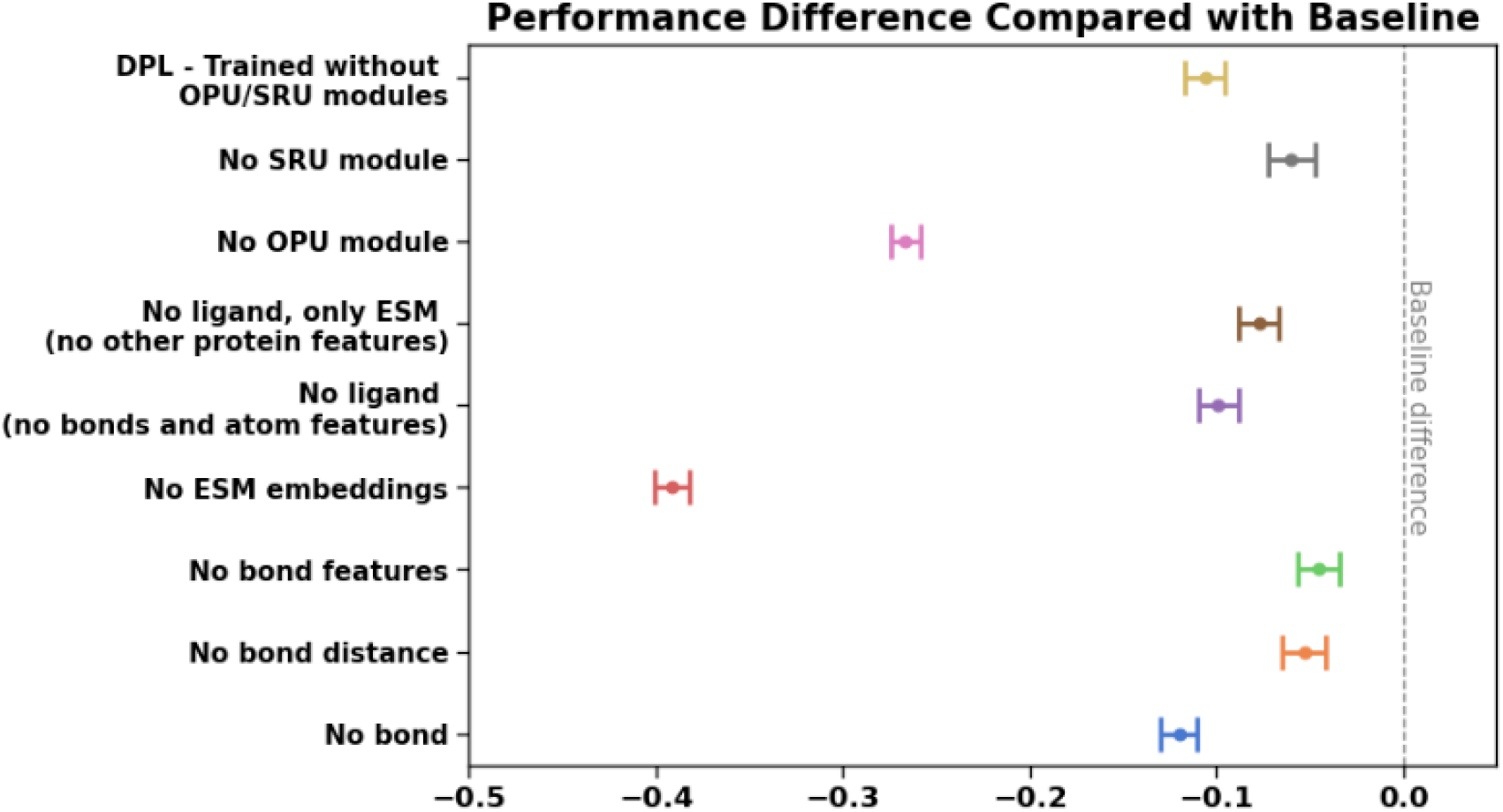
Ablation studies on ProteinReDiff’s model architecture and featurization. The dash line indicates the baseline’s average score obtained from ProteinReDiff without ablations.

We observed that our model outperformed DPL by a large margin. Incorporating just the OPU module (without the SRA module) yields better performance than DPL, indicating OPU’s ability to exchange insights between single and pair representations. Firstly, the equivariant loss function is parameterized on the structural space, making the pairwise representations from the OPU critical to that loss. Secondly, without OPU, the model performs poorly on TMScore (the bottom brown line in Figure **??**, Appendix **??**), which measures global structural preservation. Additionally, introducing SRA only without OPU hurts our model performance, suggesting the model would have been over-parameterized as the SRA updates primarily on the sequence representation. Therefore, combining both the OPU and SRA modules provides an effective approach for enhancing the representational learning of ProteinReDiff. A complete comparative assessment is presented in Table 4 and Appendix **??**.

#### Ablations on Input Featurization Methods

We conducted ablation studies to evaluate different input featurization methods, including manual feature engineering for ligands and the use of ESM-2 as a pre-trained LLM for protein featurization.

We gradually reduced ligand features, starting with ligand distance and bond information (e.g., types, ring), and even omitted the entire bond and ligand. In Figure 8, omitting bond features and distance caused less reduction in model performance than omitting the entire ligand. Ligand bond information is crucial for the model to learn the relative positions of ligand atoms and adhere to geometric constraints within the triangular update module (Section).

We observed a significant decrease in model performance when ESM embeddings were excluded (the red bar in Figure 8). The ESM features alone (the brown bar) significantly boosted performance when training without ligand data, as these embeddings are enriched with protein evolutionary and biophysical information needed for both single and pair representations. Other protein features, such as position encodings and amino acid types, provided slight improvements, though they were minimal. However, excluding ligand information led to a reduction in model performance compared to the baseline, as the model relies on learning the overall structure of the complexes.

Therefore, using pre-trained featurization methods, such as ESM and other protein BERT-like models, in combination with ligand input, significantly enhances model training and performance.

#### Impact of Masking Ratios

We examined ProteinReDiff’s performance with various percentages of masked amino acids, adjusting the masking ratio as a hyperparameter and retraining our model. In Figure 9, we observed consistent top performance across the metrics with masking ratios between 5% and 15%. This range is crucial for the protein redesign strategy, enhancing binding affinity while preserving the structural and functional motifs of the target protein. The 15% masking ratio achieved the best ligand binding affinity, the most important metric for capturing protein function.

**Figure 9:**
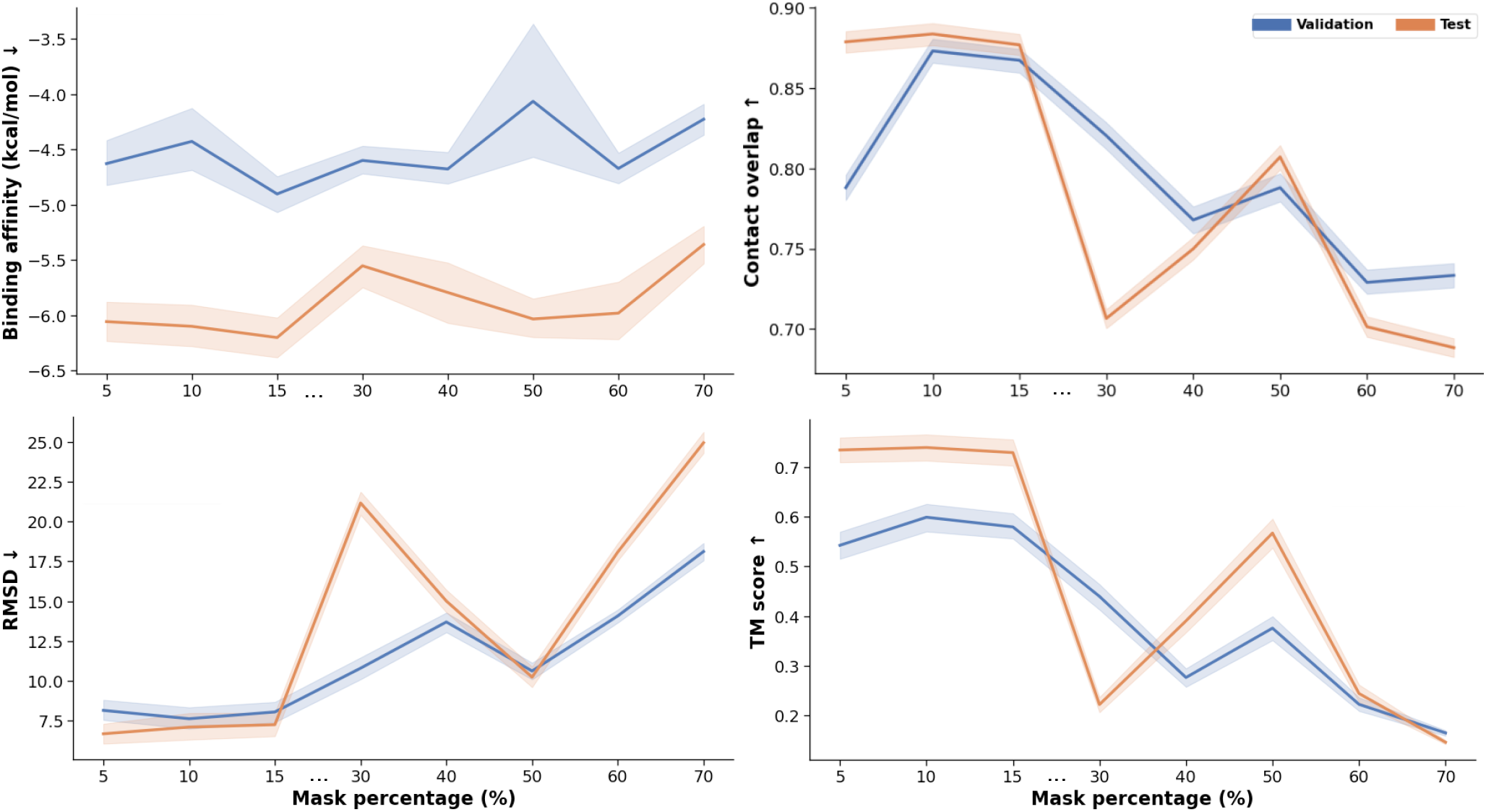
Mask ablation studies on both validation and test sets. Each of the mask ratios (5%, 10%, 15%, 30%, 40%, 50%, 60%, 70%) is a hyperparameter and represented by a model. The performances of the masked models are evaluated for all metrics. The arrows on y-axes show directions of better performance.

Interestingly, we noticed performance spikes for 50% masking in contact overlap and TMscore. This is because applying stochastic masks allows the model to learn representations with varied masking from 0 up to the set ratio. Although the 50% masking does not surpass the 15% masking’s performance, the improvement in the high masking regime demonstrates the robustness of our training scheme.

Overall, this investigation highlights the optimal level of sequence masking needed to enhance ligand binding affinity, sequence diversity, and structural preservation. It also rein-forces training strategies for protein redesign as shown on the Discussion section ().

## Conclusions

This study introduces ProteinReDiff, a computational framework developed to redesign ligand-binding proteins. By utilizing advanced techniques inspired by Equivariant Diffusion-Based Generative Models and the attention mechanism from AlphaFold2, ProteinReDiff demonstrates its ability to enhance complex protein-ligand interactions. Our model excels in optimizing ligand binding affinity based solely on initial protein sequences and ligand SMILES strings, bypassing the need for detailed structural data. Experimental validations highlight ProteinReDiff’s capability to improve ligand binding affinity while preserving essential sequence diversity and structural integrity. These findings open new possibilities for protein-ligand complex modeling, indicating significant potential for ProteinReDiff in various biotechnological and pharmaceutical applications.

## Appendix

### Appendix A: Evaluating Protein-Ligand Complex Representation

#### Evaluation Methodology

In the continuation of our study’s exploration of protein-ligand complex representations, we extended the use of the PDBBind v2020 dataset, previously detailed in our training process, to evaluate the effectiveness of embeddings generated by ProteinReDiff. Employing these embeddings as input features, we trained a Gaussian Process (GP) model aimed at predicting ligand binding affinity. The choice of a GP model recognized for its probabilistic nature and adaptability to the nuanced, uncertain dynamics of biological interactions, was pivotal in assessing how well our embeddings encapsulate predictive information about protein-ligand interactions.

#### Results and discussion

The evaluation of our ProteinReDiff model on the PDBBind v2020 dataset demonstrates competitive results in predicting ligand binding affinity using protein-ligand complex representations, as evidenced in Table 5. It’s important to note that this experiment aimed to verify the effectiveness of our protein-ligand complex representation rather than to fine-tune the model for this specific task. Consequently, while our results are promising and competitive with specialized studies focused solely on ligand binding affinity prediction, the primary goal was to validate the representation’s capability within the protein redesign framework of ProteinReDiff. This underscores the model’s utility in guiding the redesign process effectively, affirming the robustness and applicability of our protein-ligand complex representation strategy.

**Table 5:**
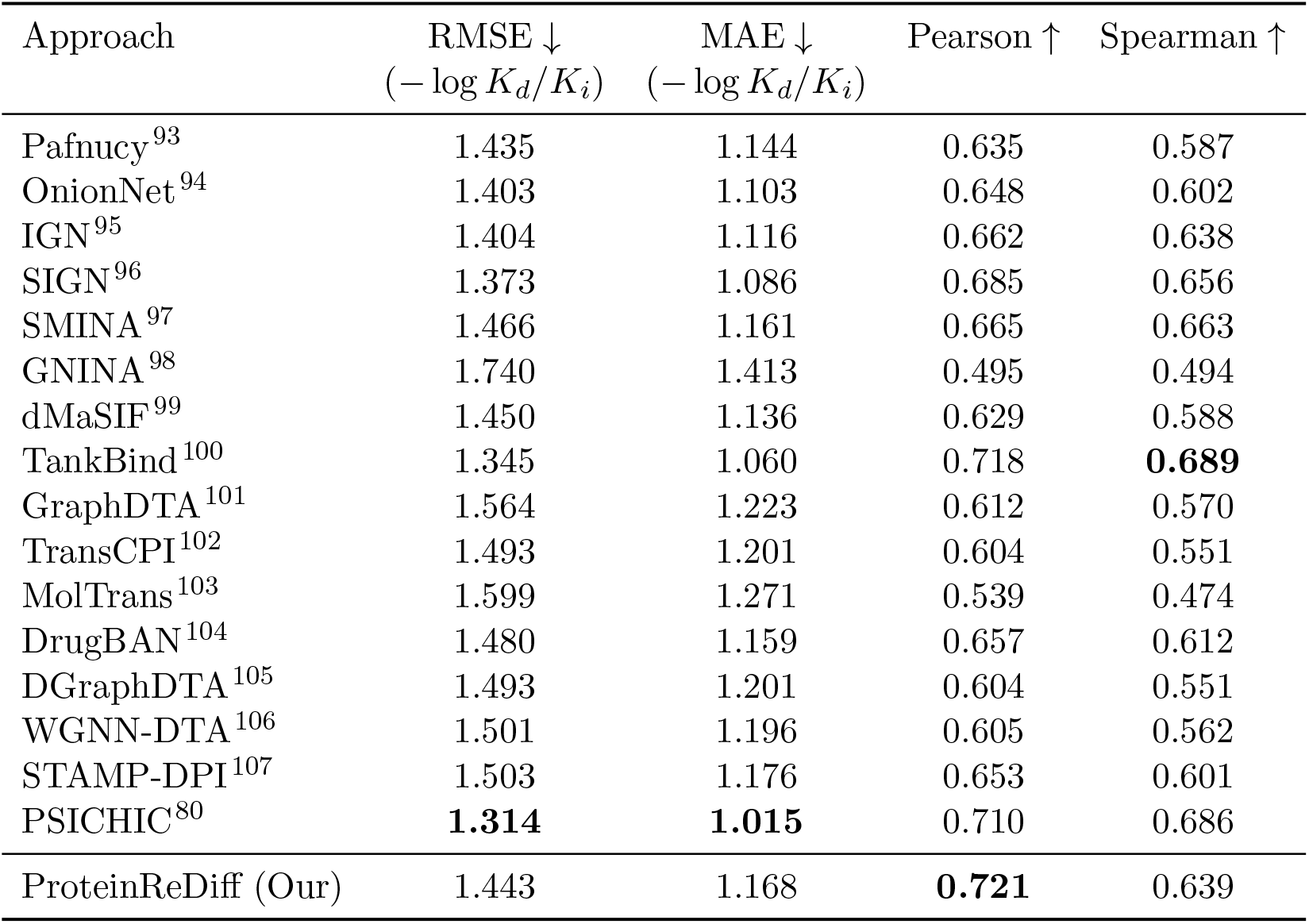
Experimental results of ligand binding affinity prediction task on PDBBind v2020 dataset.

| Approach | RMSE ↓<br>($-\log K_d/K_i$) | MAE ↓<br>($-\log K_d/K_i$) | Pearson ↑ | Spearman ↑ |
| --- | --- | --- | --- | --- |
| Pafnucy<sup>93</sup> | 1.435 | 1.144 | 0.635 | 0.587 |
| OnionNet<sup>94</sup> | 1.403 | 1.103 | 0.648 | 0.602 |
| IGN<sup>95</sup> | 1.404 | 1.116 | 0.662 | 0.638 |
| SIGN<sup>96</sup> | 1.373 | 1.086 | 0.685 | 0.656 |
| SMINA<sup>97</sup> | 1.466 | 1.161 | 0.665 | 0.663 |
| GNINA<sup>98</sup> | 1.740 | 1.413 | 0.495 | 0.494 |
| dMaSIF<sup>99</sup> | 1.450 | 1.136 | 0.629 | 0.588 |
| TankBind<sup>100</sup> | 1.345 | 1.060 | 0.718 | **0.689** |
| GraphDTA<sup>101</sup> | 1.564 | 1.223 | 0.612 | 0.570 |
| TransCPI<sup>102</sup> | 1.493 | 1.201 | 0.604 | 0.551 |
| MolTrans<sup>103</sup> | 1.599 | 1.271 | 0.539 | 0.474 |
| DrugBAN<sup>104</sup> | 1.480 | 1.159 | 0.657 | 0.612 |
| DGraphDTA<sup>105</sup> | 1.493 | 1.201 | 0.604 | 0.551 |
| WGNN-DTA<sup>106</sup> | 1.501 | 1.196 | 0.605 | 0.562 |
| STAMP-DPI<sup>107</sup> | 1.503 | 1.176 | 0.653 | 0.601 |
| PSICHIC<sup>80</sup> | **1.314** | **1.015** | 0.710 | 0.686 |
| ProteinReDiff (Our) | 1.443 | 1.168 | **0.721** | 0.639 |

